# Disparity analyses are robust to ancestral state estimation uncertainty

**DOI:** 10.64898/2026.04.21.719166

**Authors:** Caleb Scutt, Natalie Cooper, Gavin H. Thomas, Thomas Guillerme

## Abstract

Morphological trait datasets and phylogenies are routinely paired to investigate macroevolutionary patterns during disparity analyses. However, incomplete fossil sampling can distort disparity estimates, obscuring true evolutionary signals. Ancestral state estimation can be used for both continuous and discrete traits to extend these analyses beyond incomplete fossil data, such as investigations into disparity through time. However, when ancestral state estimation occurs in the disparity pipeline, and the inevitable uncertainty in these estimates, complicate their integration. Determining the most robust workflow for integrating ancestral state estimation in disparity analyses remains a critical methodological challenge. Using simulations to attain a ground-truth disparity value, we evaluated different approaches to performing ancestral state estimation and incorporating uncertainty across varying continuous and discrete trait models, fossil sampling densities and disparity metrics. Ancestral state estimation generally improved recovery of true disparity relative to tip-only analyses, though the optimal approach depended on the interaction between trait model and fossil sampling density. For continuous traits, probabilistic approaches were most accurate, but were sensitive to model misspecification under low fossil sampling density. For discrete traits, pre-ordination methods were most reliable and probabilistic approaches outperformed point estimates under low sampling, while point estimates became increasingly accurate as sampling density increased. Fossil sampling density was a stronger predictor of disparity accuracy than estimation method choice, underscoring that methodologies are only as powerful as the data provided. Our findings offer a practical decision framework for selecting the most appropriate workflow given the sampling density and trait characteristics of a dataset.

## 1 Introduction

Morphological diversity, hereafter termed disparity, is widely used for investigating macroevolutionary dynamics (Foote 1997; Wills 2001; Guillerme et al. 2020a). Disparity can be described as the distribution or spread of taxa within a multidimensional trait space defined by morphological similarity (Erwin 2007) and can be quantified using both continuous (García-Navas et al. 2017; Gray et al. 2019; Friedman et al. 2020) and discrete characters (Foote 1994; Lloyd 2016). Morphological trait datasets are routinely integrated with phylogenies when examining disparity across clades or time. These analyses can be used to investigate selectivity across mass extinctions (Bapst et al. 2012; Cole and Hopkins 2021; Liu et al. 2024), reveal patterns in morphological convergence (Muschick et al. 2012; Stayton 2015, 2024) or characterise the tempo and mode of morphological diversification during adaptive radiations (Harmon et al. 2003; Mahler 2010; Hughes et al. 2013). However, such analyses are inevitably limited by the incompleteness of the fossil record; only a biased subset of past diversity is preserved, leaving unsampled ghost lineages and obscuring large portions of a clade’s evolutionary history (Flannery Sutherland et al. 2019; Smith et al. 2021). These missing data can result in both under- and over-estimations of disparity (Mitchell 2015; Smith et al. 2023).

One way in which unsampled lineages can be accounted for is through the use of ancestral state estimation. Ancestral state estimation is the method of estimating trait value(s) in a set of common ancestors, i.e., nodes on a phylogenetic tree, given a set of tip data (Schluter et al. 1997; Cunningham et al. 1998; Revell 2025). These estimated states can be combined with the observed tip data to generate a more complete trait space, filling the gaps left by an incomplete fossil record and, in theory, improving our ability to recover the ‘true disparity’ of a group (Sidlauskas 2008; Brusatte et al. 2011; Lloyd 2018). True disparity in this context can be defined as the total trait space occupied by all lineages throughout a clade’s history, measured by either volume or density (Guillerme et al. 2020b). Ultimately, the inclusion of ancestral states allows disparity through time analyses to be explored with higher temporal resolution (Brusatte et al. 2011; Halliday and Goswami 2016; Guillerme 2018).

Despite the aforementioned advantages, the use of ancestral state estimation within disparity analyses has often been neglected. The hesitancy to incorporate ancestral state estimation has likely stemmed from the notion that estimates are inherently uncertain; root-ward estimates in particular are often associated with large confidence intervals or inconclusive scaled likelihoods (Schluter et al. 1997; Losos 1999; Gascuel and Steel 2014). While the propagation of uncertainty during comparative analyses is encouraged (Silvestro et al. 2015; Boyko and O’Meara 2024), most computational implementations of ancestral state estimation return point estimates, because there is little guidance on how to correctly treat ancestral state estimation uncertainty downstream when constructing the trait space. These point estimates are typically averages from confidence intervals for continuous characters or collapsed states returned from arbitrary thresholds for discrete characters (Smith and Donoghue 2022; Casali et al. 2023). By ignoring ancestral state estimation uncertainty, point estimates can lead to overconfidence in results and incorrect inferences in downstream analyses, all while obscuring potentially accurate signals (Ackerly et al. 2006; Vanderpoorten and Goffinet 2006).

Another key issue with ancestral state estimation is its sensitivity to model misspecification. Likelihood-based approaches to ancestral state estimation require both a tree topology and a specified model of character evolution. Continuous character estimations are typically modelled under Brownian motion (BM; but see Elliot and Mooers 2014; Castiglione et al. 2020; Meade and Pagel 2022), which describes a directionless, stochastic process where the trait divergence between related taxa is proportional to their shared evolutionary history and the rate of evolution (Felsenstein 1985; Schluter et al. 1997; O’Meara et al. 2006). However, deviations from BM in the trait data, such as directional trends or stabilising selection, inevitably returns biased estimates (Webster and Purvis 2002; Finarelli and Flynn 2006; Albert et al. 2009; Finarelli and Goswami 2013). Model-related issues can also arise for discrete traits, which are estimated using M*k* (Lewis 2001; Pagel 1994, 1999) or *SSE models (Ng and Smith 2014; Holland et al. 2020). Estimation accuracy of discrete states appears prone to model over-parameterisation and sensitive to fast transition rates (Mooers and Schluter 1999; Keating 2023). Furthermore, discrete ancestral state estimation faces a paradoxical trade-off where a slow rate of evolution allows for an accurate estimation of the root state, but provides little information for calculating transition rates (Gascuel and Steel 2020).

Given that uncertainty is often high in ancestral state estimation, how then can estimates be improved? Ancestral state estimation is statistically inconsistent, meaning that even if the sample size of extant tips were infinite, estimates are not expected to converge on the true value (Ané 2008). The inclusion of fossils can improve estimations by breaking long branch lengths, reducing distance between tips and internal nodes (Finarelli and Flynn 2006; Finarelli and Goswami 2013; Joy et al. 2016; Goswami and Clavel 2025). Numerous simulation and empirical based studies have shown that ancestral state estimations based on extant-taxa only are extremely biased (Webster and Purvis 2002; Finarelli and Flynn 2006; Albert et al. 2009; Betancur-R et al. 2015; Royer-Carenzi and Didier 2016; Monson et al. 2022; Wisniewski et al. 2022; Keating 2023), though this bias can be driven by model misspecification rather than missing data alone (Goldberg and Igić 2008; Puttick and Thomas 2015). Similar biases have been observed in extant-only disparity analyses (Mitchell 2015). Despite this, many studies are forced to use extant-only trees for both ancestral state estimation (Jetz and Pyron 2018; Amador and Giannini 2021) and disparity (Harmon et al. 2010; Baker et al. 2015; Smith and Donoghue 2022) due to a lack of viable fossil data.

Numerous studies have assessed the raw estimation error of ancestral state estimation approaches under different models of evolution, methods of estimation and fossil sampling density (Elliot and Mooers 2014; Royer-Carenzi and Didier 2016; Reyes et al. 2018; Holland et al. 2020; Keating 2023; Boyko et al. 2026). However, how trait estimation accuracy is propagated into the recovery of an accurate trait space is less understood, despite the continued use of ancestral state estimation in disparity analyses. One methodological debate concerns whether ancestral state estimation should be conducted before or after trait space ordination. Previous work has found that carrying out ancestral state estimation on the axes of an ordination matrix rather than raw characters can distort the distribution of points in a trait space markedly, introducing unwanted phylogenetic signal and low levels of convergence (Lloyd 2018). However, this study was restricted to a single empirical dataset and discrete traits. Thus, we lack a comprehensive understanding of how estimation approaches, trait data and fossil sampling density interact to affect our ability to recover accurate patterns in trait space and therefore disparity.

Here we evaluate the ability of different ancestral state estimation approaches to recover true disparity values across different evolutionary models and fossil sampling densities using a simulation framework. Simulations offer a robust approach for benchmarking methodologies because a ground-truth dataset can be generated, so trait values and the resulting disparity value are all known *a priori* (Lotterhos et al. 2022). Different methodological choices and estimation scenarios can then be evaluated against the true disparity value. Specifically, we directly compare pre-versus post-ordination ancestral state estimation approaches, as well as point-estimate versus probabilistic treatments of uncertainty. We also investigate how estimation error varies under different models of continuous (Brownian motion with and without trend and Ornstein-Uhlenbeck) and discrete traits (varying transition rates), fossil sampling densities (100%, 50%, 15%, 5% and 0% of extinct lineages) and disparity metrics examining size and density. This integrative approach provides a comprehensive evaluation of when ancestral state estimation improves disparity estimation, when it introduces bias, and how uncertainty should be treated when working with an incomplete fossil record.

## 2 Materials and Methods

### 2.1 Simulation Framework

#### 2.1.1 Phylogenies

All simulations were generated using treats in R v.4.5.2 (Guillerme 2024; R Core Team 2025). We simulated birth-death trees with the speciation (*λ*) parameter set to 1.0, and extinction (*µ*) set to 0.9 (Figure 1 Step 1). Fixing *λ* = 1.0 provides a convenient relative timescale, because only the ratio of extinction to speciation affects the shape and balance of simulated trees. These parameters correspond to a high turnover regime, consistent with the upper-empirical estimates from diversification studies of animal groups (Scholl and Wiens 2016; Pires et al. 2018; Henao Diaz et al. 2019; Morlon et al. 2024). This fraction of extinction (*µ* = 0.9) produces trees that match our expectation of substantial extinct diversity based on the fossil record.

**Figure 1:**
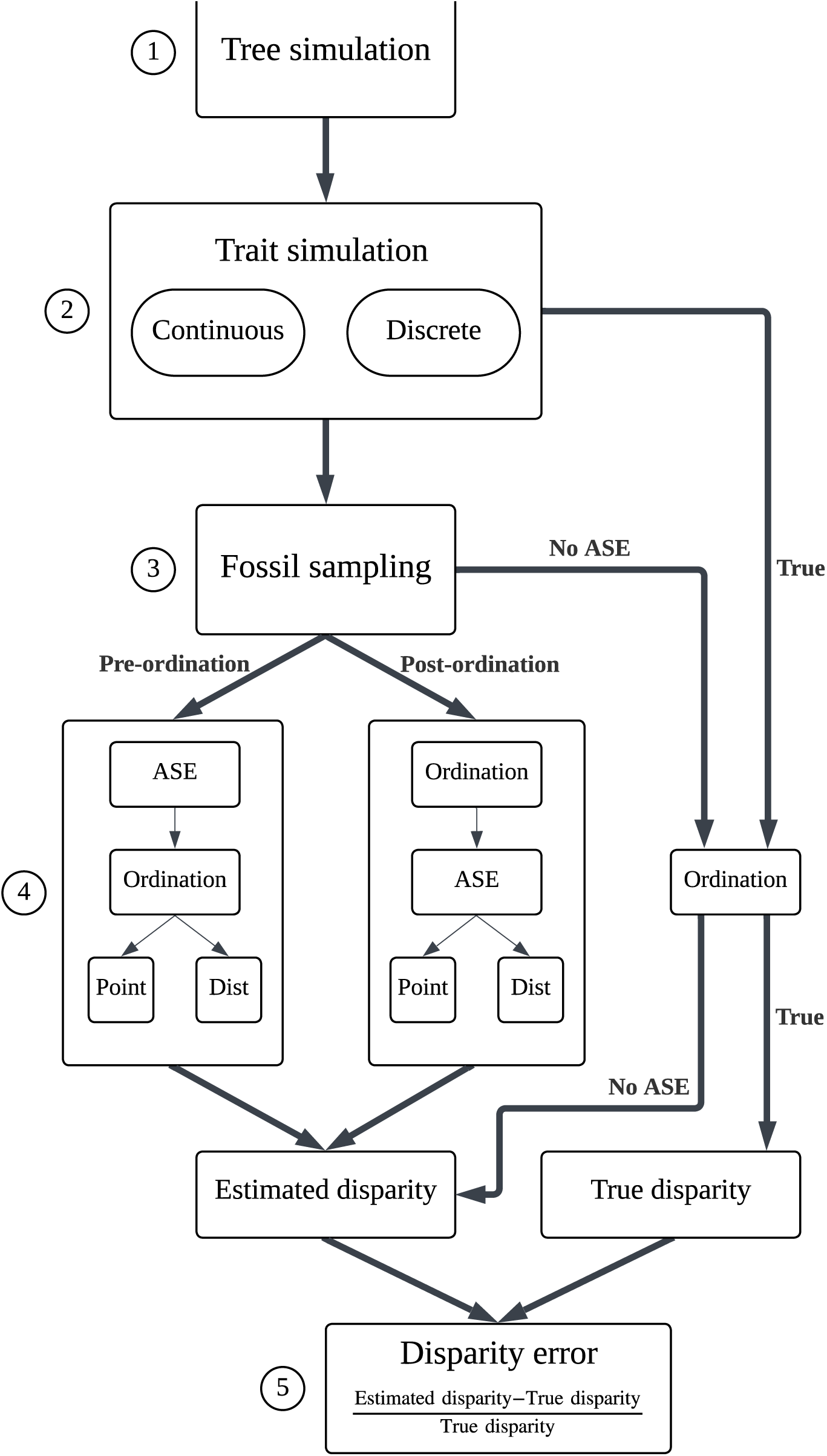
Overall simulation pipeline, highlighting the five different estimation routes (pre-ordination point, pre-ordination distribution, post-ordination point, post-ordination distribution and no ancestral state estimation) as well as the true route, from which ground-truth disparity value is calculated from the untreated simulation.

We generated three sets of 100 birth-death trees with these parameters until the simulations reached 50, 100 or 150 extant taxa, with maximum tree size of 500, 1000 and 1500 tips, respectively (Figure S1). The randomness of the birth-death model meant some simulated trees contained only extinct taxa, so we re-ran each individual simulation until we obtained the desired number of extant taxa (50, 100 or 150). This meant that although the parameters were set to *λ* = 1 and *µ* = 0.9, the effective parameters slightly deviated from these input values (Figure S2). Mean net diversification across all simulated trees was 0.30 *±* 0.09.

Trees were pruned to extract the crown clade of the tree for downstream analyses. This ensured that all simulated fossil sampling densities shared an identical root node and tree height, providing a consistent framework for downstream analyses. Any tip included in the analyses was a descendant of the most basal node considered, i.e., the MRCA of extant tips, ensuring that all ordinations and disparity values were calculated within a consistent clade. This establishes a robust assessment of the interaction between fossil sampling and ancestral state estimation.

#### 2.1.2 Trait simulations

For each phylogeny, we simulated continuous and discrete trait data separately (Figure 1 step 2). Continuous trait data were simulated under variations of Brownian motion (BM) and Ornstein-Uhlenbeck (OU) models (Table 1). We simulated BM with variance = 0.25. We also simulated BM with a trend parameter of 0.3, where the ancestral trait value increased incrementally proportional to branch length. OU trait data was simulated with the *α* parameter set based on phylogenetic half-life to account for differing tree heights across simulations (Gearty et al. 2024).

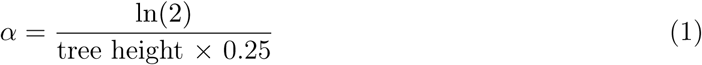

We simulated 20 independent characters for each model (BM, BM + trend and OU) across each phylogeny.

Discrete trait evolution was simulated using a discrete-time Markov process, where the probability of state transition along each branch was proportional to the product of the transition probability and branch length. We simulated discrete traits across three rate categories, representing fast (0.9), medium (0.1) and slow (0.01) transition rates (Table 1). These rates were assigned by generating equal-rates (ER) transition probability matrices. For each category, 85 binary characters and 15 three-state characters were simulated, following a typical scoring ratio found in empirical data (Guillerme and Cooper 2016; Casali et al. 2023).

**Table 1:**
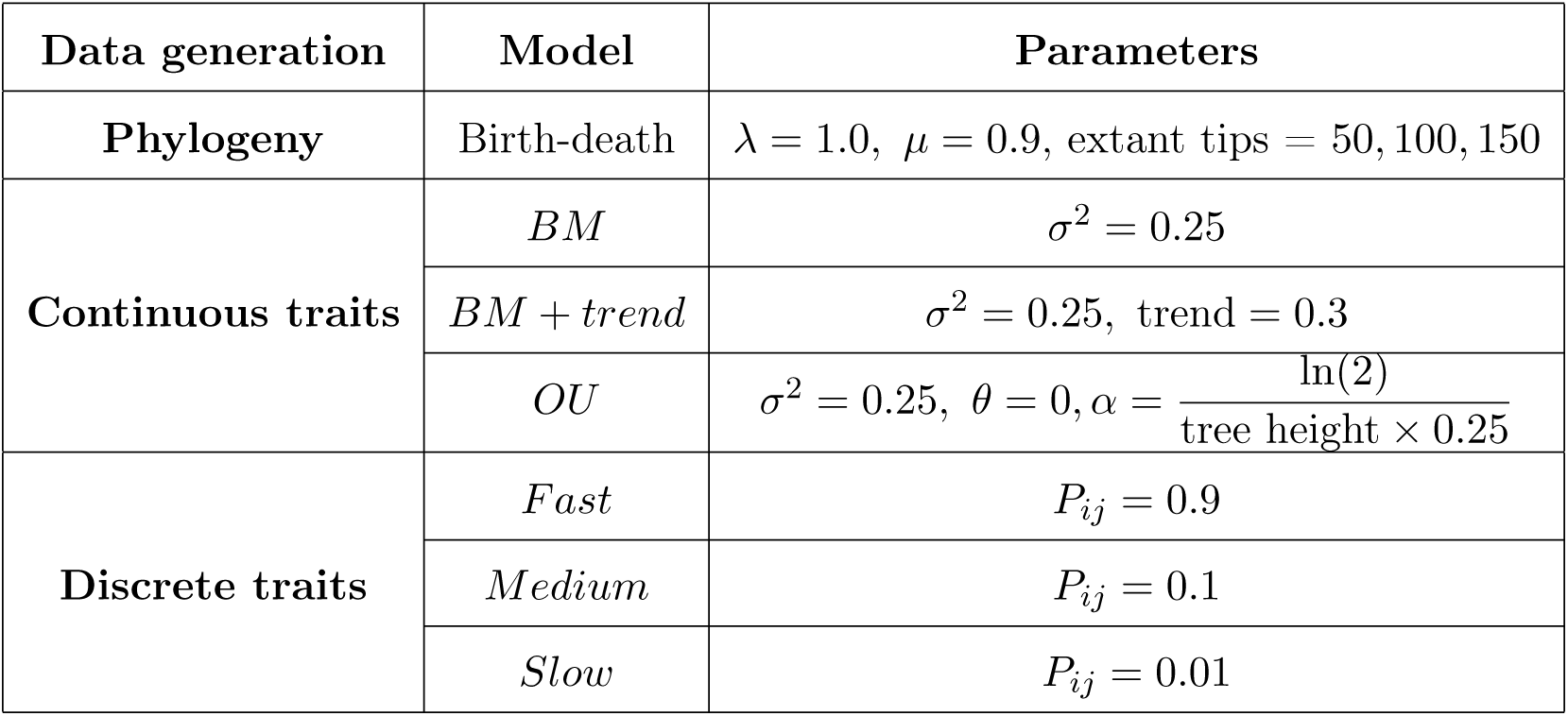
Simulation models and parameters used for phylogeny generation and trait evolution. For continuous characters, the root state was always set to 0. *λ* = speciation; *µ* = extinction; *P_ij_* = transition rate; *σ*^2^ = evolutionary variance; *θ* = optimum, *α* = rate of adaptation toward the optimum.

### 2.2 Fossil sampling simulation

To simulate different fossil sampling densities, extinct tips were either retained (‘fossilised’) or pruned using a Bernoulli sampling method at five different probabilities: 0%, 5%, 15%, 50% and 100% (Figure 1: Step 3). This method means each extinct tip had an equal probability of being ‘fossilised’ at each sampling level, irrespective of the total tree size. The corresponding trait matrices were subset to match the tips remaining in each down-sampled tree.

### 2.3 Ancestral state estimation (ASE)

We estimated ancestral state on the down-sampled matrices using `dispRity::multi.ace` (Guillerme 2018), which is a wrapper for `ape::ace` (Paradis and Schliep 2019). We used four combinations of two approaches to ancestral state estimation to evaluate each method’s accuracy in recovering the true disparity signal (Figure 1: Step 4). These were then compared to a null model bypassing ancestral state estimation.

#### 2.3.1 Pre-ordination estimation

Ancestral state estimation of trait data can be carried out at two points either side of the ordination step in the pipeline (Figure 1: Step 4; Lloyd 2018). Pre-ordination ancestral state estimation estimation involves the estimation of the raw trait values for each node. We performed pre-ordination estimation of continuous characters under a BM model, where ancestral state values lie within a multivariate normal distribution that is estimated given the observed tip data and the shared evolutionary history among taxa (Schluter et al. 1997; Rohlf 2001). The width of the estimated confidence interval returned reflects the uncertainty in continuous ancestral estimates. This width typically increases with node depth and decreases with the number of informative descendant lineages.

Pre-ordination estimation of discrete characters involves estimating the marginal likelihoods for each discrete state at each node, based on a maximum likelihood derived estimate of the M*k* transition matrix and the tree topology given (Pagel 1999; Yang 2006). Given that the transition rates were set *a priori* during simulations, the equal-rates (ER) model was used to avoid model misspecification as a confounding factor on estimation accuracy (Keating 2023).

We constructed trait spaces for disparity estimation by ordinating the matrices of combined tip and ancestral states. Continuous data were ordinated using principal components analysis (PCA; `stats::prcomp` function in R). Because PCA requires raw continuous data to calculate covariance, discrete characters were instead converted to distance matrices using the Maximum Observed Rescaled Distance metric (MORD) (Lloyd 2016) and ordinated using principal coordinates analyses (PCoA) (`stats::cmdscale` function in R) with the Cailliez correction to deal with negative eigenvalues (Cailliez 1983).

#### 2.3.2 Post-ordination estimation

Post-ordination estimation calculates the continuous PC scores across each axis of an ordination matrix for each ancestral node. For post-ordination ancestral state estimation, PCA and PCoA were applied prior to ancestral state estimation to the down-sampled continuous and discrete character matrices, respectively, to generate a set of tip-only trait spaces. Ancestral node states were then estimated under a Brownian motion model across all PC axes, as detailed in subsubsection 2.3.1.

#### 2.3.3 Uncertainty treatment

For each estimation method, we also compared point estimate and probabilistic approaches. The point estimate approach returns a fixed single value for each estimate. For continuous characters, the point estimate returned by `ape::ace` is the single maximum likelihood estimate of the ancestral state. For discrete characters, we used a strict majority rule where the state with the highest scaled likelihood was selected at each node for each character. In scenarios where state likelihoods were equal, one state was chosen at random.

The probabilistic approach differs in that uncertainty was explicitly accounted for by sampling multiple estimates rather than a single point. For continuous characters, we sampled each state across 100 matrices, with each estimate value lying within a normal distribution centered on the single point estimate and spread across the 95% confidence interval. Each discrete character state at each node was also sampled across 100 matrices according to their scaled likelihoods. In a scenario where there is uncertainty in a binary character at a specific node, e.g., 0.55 for state 0 and 0.45 for state 1, this distribution approach results in a 55% probability of sampling state 0 in each matrix for the respective node (and 45% chance of sampling state 1). Both procedures were implemented using the `sample = 100` argument in `dispRity::multi.ace` (Guillerme 2018).

### 2.4 Disparity analyses

Our aim for this study was to understand how well ancestral state estimation can recover true signals in size and density in trait space occupancy (Guillerme et al. 2020b). Size-based metrics describe the volume of trait space the clade occupies (Guillerme et al. 2020b). To examine how well the size of the trait space was recovered, we calculated the sum of quantiles. The sum of quantiles calculates the range on each PC axis for the 95% of the data and then sums these ranges across all axes. This metric has similar properties to the often used sum of ranges metric by describing the total trait space exploited (Foote 1992), but with less sensitivity to outliers and sample size. For density, we calculated the sum of variances (Ciampaglio et al. 2001; Wills 2001) and the mean pairwise distance (Ciampaglio et al. 2001). To quantify methodological performance, we calculated these disparity metrics for the ground-truth simulated trait spaces, the tip-only trait spaces, and the spaces generated by each ancestral state estimation approach so that relative disparity errors could be calculated (Figure 1 step 5).

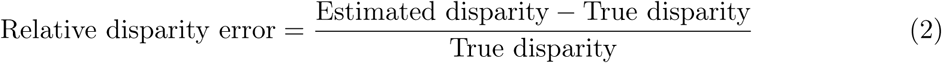

### 2.5 Statistical tests

The relative disparity error values were converted to absolute errors to measure the magnitude of deviation from the true disparity, regardless of direction. Due to the zero-inflated and right-skewed nature of the error distribution, we applied a log-transformation. Consequently, the more negative the log-error, the more accurate (closer to *log*(0)) the original estimation. Given the imbalance in sample sizes between the different uncertainty treatments (i.e., distribution approaches return 100 disparity values per estimate, whereas point estimates return one), we aggregated the distribution outputs by taking the median prior to analyses. We also performed analyses where the distribution was retained, but each individual point returned by distributions was weighted 1/100th of the single point estimate. Results were consistent between both approaches (see supplements).

We investigated the drivers of disparity estimation accuracy with linear-mixed models (LMM) using the `lme4` R package (Bates et al. 2015; Figure S4, S5) The log-transformed relative error was treated as the response variable. Based on our experimental design, we fitted a four-way interaction between the estimation method, model of evolution, fossil sampling density and disparity metric. We also included tree size as a further non-interactive fixed effect. To account for the variation with each simulation run, each tree replicate was included as a random intercept.

We carried out *post-hoc* pairwise comparisons using estimated marginal means, adjusted using Tukey’s honestly significant difference (HSD) (Lenth and Piaskowski 2025).

## 3 Results

The results reported herein are from the aggregated linear mixed-effects models (LMMs). Results from the alternative weighted LMMs were consistent with these findings (Figure S6, S7).

### 3.1 Continuous traits

#### 3.1.1 Estimation methods

For continuous characters, the estimation method had a significant effect on disparity accuracy (*F*_4,66976_ = 358, *p <* .001; Figure 2). Notably, using ancestral state estimation methods improved estimates of disparity compared to excluding ancestral states when averaged over all scenarios. *Post-hoc* pairwise comparisons of estimated marginal means (EMM) revealed both pre-and post-ordination distributions of ancestral state estimations returned the lowest log disparity errors (Table S1). In contrast, point estimates performed worse overall than bypassing ancestral state estimation altogether.

**Figure 2:**
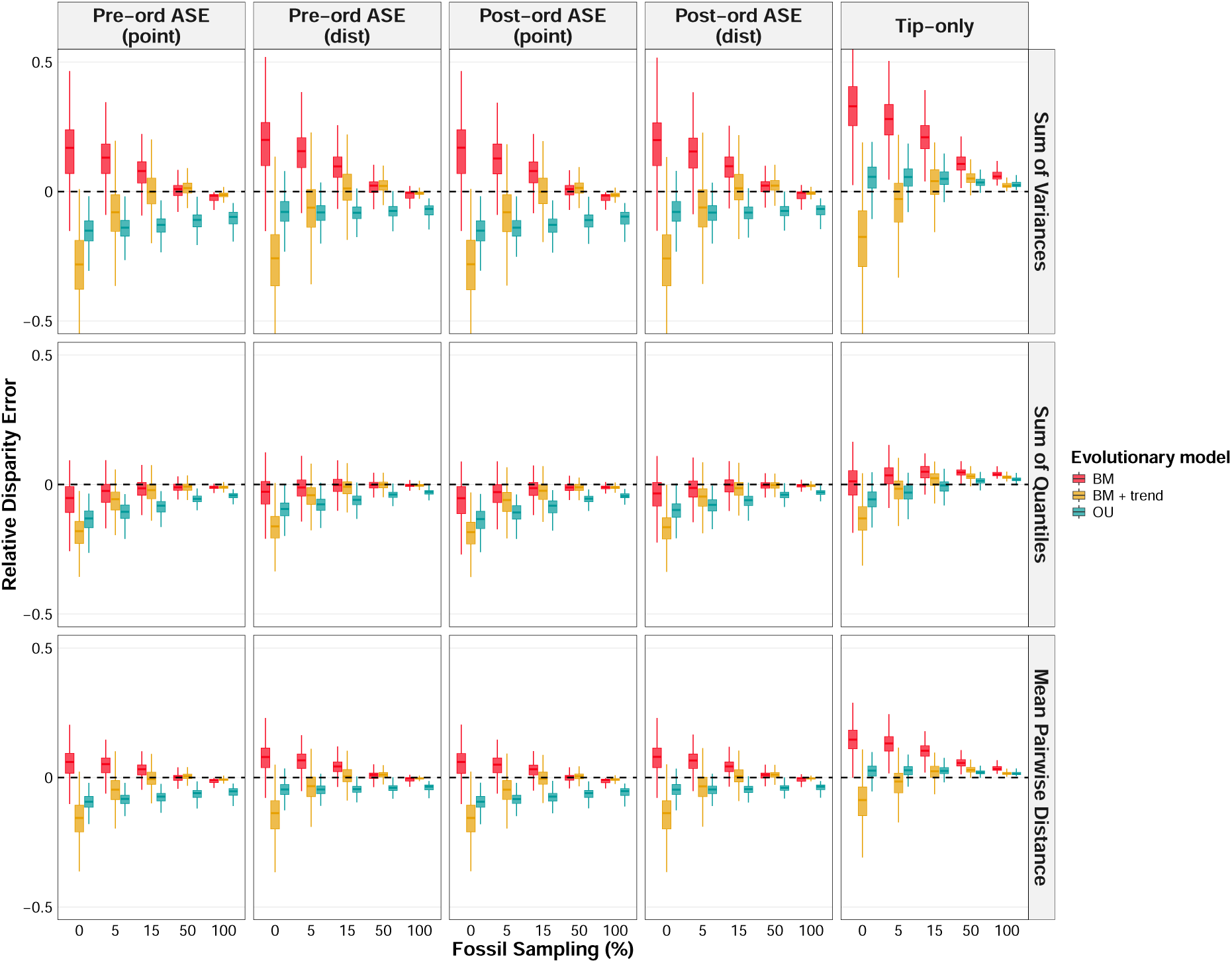
Relative disparity error returned from pre-ordination point ancestral state estimation (ASE), pre-ordination distribution ASE, post-ordination point ASE, post-ordination distribution ASE and no ASE. These disparity errors were calculated across three evolutionary models (Brownian motion (BM), BM + trend and OU), five simulated fossil sampling densities (100%, 50%, 15%, 5% and 0%) and three disparity metrics (Sum of variances, sum of quantiles and mean pairwise distance). Values closest to 0 (dotted horizontal line) were most accurate; negative errors are under-estimates and positive are over-estimates.

#### 3.1.2 Interaction effects

The pattern that ancestral state estimation methods improved estimates of disparity was context-specific (Figure 3). Fossil sampling was the strongest driver of recovery of an accurate disparity signal for continuous traits (*F*_4,66976_ = 9169, *p <* .001), as each increase in fossil sampling density resulted in a significant reduction in disparity error. Similarly, the evolutionary model (*F*_2,66976_ = 2956, *p <* .001) and disparity metric (*F*_2,66976_ = 5028, *p <* .001) were also strong drivers of variation in error. Specifically, disparity outputted from the BM models and sum of quantiles metric were associated with better disparity estimates, whereas the OU model and sum of variances were the least accurate. Tree size had a minor effect in comparison (*F* = 43_2,297_, *p <* .001). The three-way interaction between estimation method, fossil sampling and evolutionary model was significant (*F*_32,66976_ = 60, *p <* .001) and the *post-hoc* EMM can be used to identify the optimal method for models at each fossil sampling density (Table S2). For Brownian motion (BM) at 100% fossil sampling, the distribution estimations were most accurate. At 50% and below, methods of ancestral state estimation generally performed equally well and significantly better than excluding ancestral states. Under BM + trend, a similar pattern was followed at high fossil sampling densities, yet when fossil sampling dropped below 15% methods of ancestral state estimation performed worse than excluding ancestral states. In contrast, under an OU model the exclusion of ancestral states was the best approach across all fossil sampling densities. While the disparity metric had a large effect on the overall estimation accuracy, it was a relatively lower driver of variation regarding its interaction with the estimation method (*F*_8,66976_ = 7, *p <* .001). Therefore, while estimation accuracy differed between metrics, the hierarchy between optimal methods was generally consistent across metrics.

**Figure 3:**
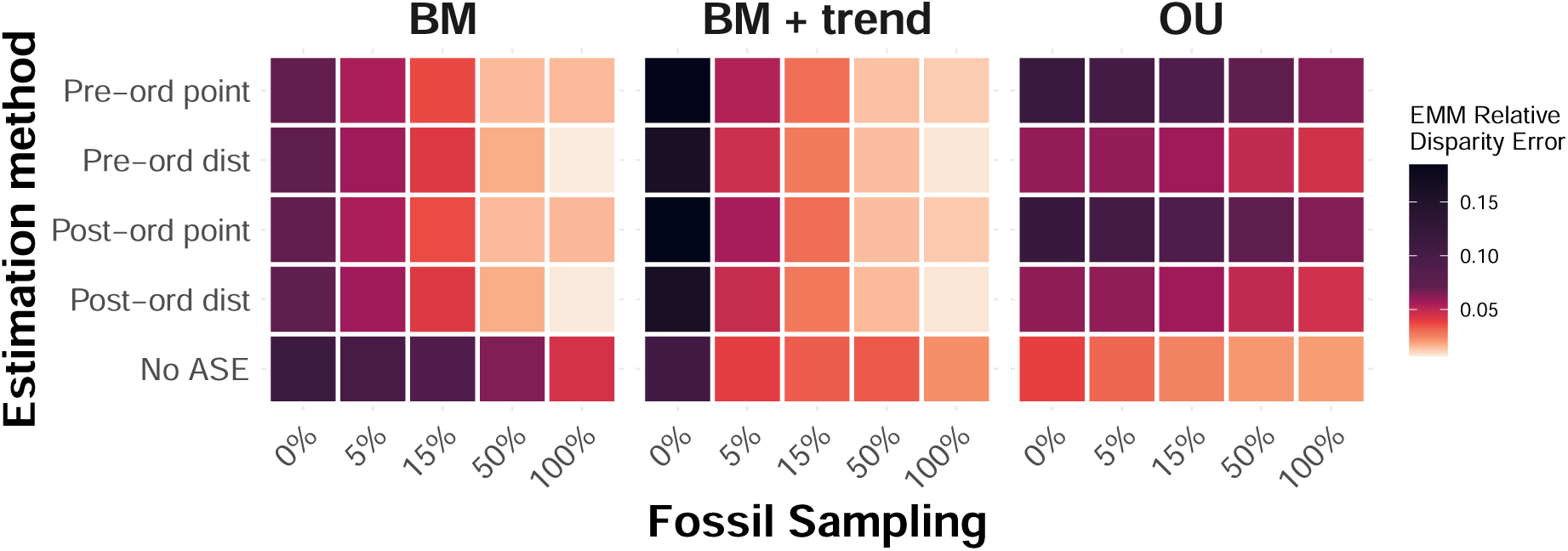
Heatmap presenting the continuous estimated marginal mean (EMM) relative disparity errors for every combination of estimation method, evolutionary model and simulated fossil sampling density. EMM were calculated from a linear mixed model with a three-way interaction term, averaged across disparity metrics.

### 3.2 Discrete traits

#### 3.2.1 Estimation methods

For discrete characters, the estimation method also had a significant effect on disparity accuracy (*F*_4,66976_ = 10767, *p <* .001; Figure 4). Similar to continuous traits, methods of ancestral state estimation consistently improved disparity estimates compared to excluding ancestral states. However, the optimal type of estimation differed. *Post-hoc* pairwise comparisons of EMM revealed that pre-ordination point estimates returned the lowest log disparity errors overall. In contrast to continuous traits, distribution methods generally performed worse than pre-ordination point estimates on average, but post-ordination point estimates performed worse than excluding ancestral states (Table S3).

**Figure 4:**
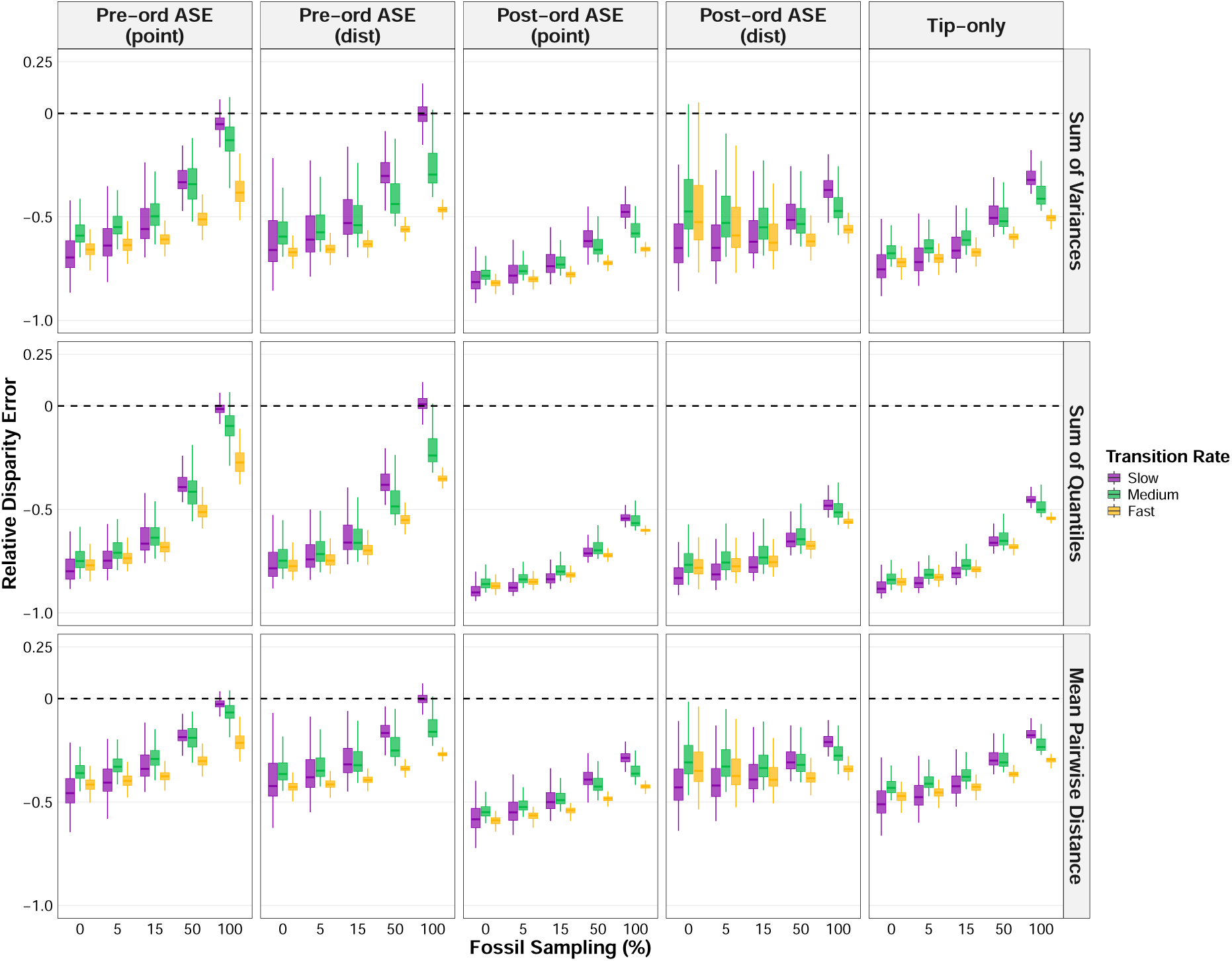
Relative disparity error returned from pre-ordination point ASE (ancestral state estimation), pre-ordination distribution ASE, post-ordination point ASE, post-ordination distribution ASE and no ASE. These disparity errors were calculated across three transition rates (fast (0.9), medium (0.1) and slow (0.01)), five simulated fossil sampling densities (100%, 50%, 15%, 5% and 0%) and three disparity metrics (Sum of variances, sum of quantiles and mean pairwise distance). Values closest to 0 (dotted horizontal line) were most accurate; negative errors are under-estimates and positive are over-estimates.

#### 3.2.2 Interaction effects

Similar to continuous traits, fossil sampling (*F*_4,66976_ = 22191, *p <* .001), transition rate (*F*_2,66976_ = 6957, *p <* .001) and metric (*F*_2,66976_ = 26174, *p <* .001) were again major drivers of variation in accuracy. As expected, increased fossil sampling and decreased transition rate led to lower disparity errors, and the mean pairwise distance was the easiest metric to recover, whereas the sum of quantiles was the hardest. The three-way interaction between estimation method, fossil sampling and transition rate explained much of the variation in the data (*F*_32,66976_ = 588, *p <* .001; Table S4). The broad pattern was that at high fossil sampling densities, pre-ordination point estimates were the best method, whereas at lower fossil sampling densities both pre- and post-ordination distributions returned the most accurate disparity values (Figure 5). This pattern was model-dependent, however, as under a slow rate the pre-ordination distribution method was the best performing method across all fossil sampling densities. At fast and medium rates under low fossil sampling (< 15%), the post-ordination distribution method returned the lowest disparity errors. Notably, methods of ancestral state estimation outperformed their exclusion across all combinations of transition rate, fossil sampling density and disparity metric. Again, while the disparity metric was a significant driver to overall accuracy, it had little effect on the hierarchy in methods set by the transition rate and fossil sampling density, signified by a weak four-way interaction (*F*_64,66976_ = 10, *p <* .001).

**Figure 5:**
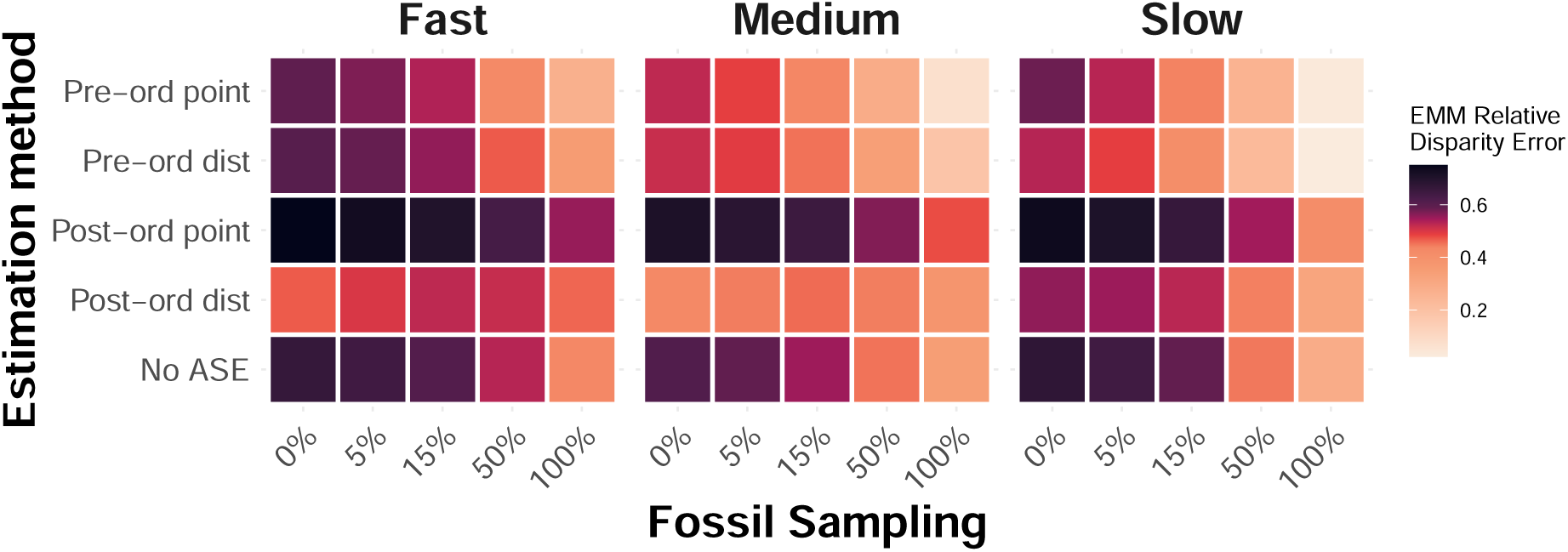
Heatmap showing the discrete estimated marginal mean (EMM) relative disparity errors for all combinations of estimation method, transition rate and simulated fossil sampling density. EMM were calculated from a linear mixed model with a three-way interaction, averaged across disparity metrics.

## 4 Discussion

Our results suggest that, generally, the inclusion of ancestral states improves the recovery of true disparity, defined here as the total region in trait space occupied by all members of a clade that have ever lived. However, we found this to be context-specific, particularly dependent on fossil sampling and model of evolution. Overall, we recommend that ancestral state estimations can consistently improve estimations of disparity derived from discrete characters, but recommend caution for continuous characters.

### 4.1 Pre- versus post-ordination estimation

For continuous characters, we found virtually no discrepancy between the effect of pre- and post-ordination estimation of ancestral states on disparity. Given that continuous traits exist within a Euclidean space, the linear transformations of both PCA and ancestral state estimation under Brownian motion (BM) mean that estimated node states exist within the same relative positions to the observed tip data for both pre- and post-ordination estimation. Note, however, that our simulated traits were uncorrelated. In the case of non-orthogonality between traits, it is likely that the position of ancestral state estimation in the disparity pipeline will have some effect, although this was not tested here.

For discrete traits, the position of ancestral state estimation relative to ordination during the disparity pipeline (Figure 1) was a key driver in accurate recovery of the true disparity. Overall, pre-ordination estimation led to more accurate disparity estimations than post-ordination estimation, a finding consistent with previous work (Lloyd 2018). Post-ordination estimation is ultimately limited by the definition of the resulting trait space. Axes of the trait space are treated as continuous traits evolving under BM, so, given the assumptions of BM, ancestral nodes are inevitably constrained within the bounds per-PC axis defined by the observed tip data (Stayton 2015; Lloyd 2018; Gerber 2019). This forces the implicit assumption that our observed trait data accounts for the extremes of morphological diversity for the clade of interest, and that past diversity is only a subset of sampled diversity. Disparity is therefore underestimated, particularly with range-based metrics such as sum of quantiles (Figure 4). Pre-ordination estimation does not encounter these same limitations. A discrete trait matrix exists within a hypercube, where each vertex corresponds to a unique combination of traits. When ancestral states are estimated prior to ordination, there is no constraint preventing an ancestor from occupying a legitimate and theoretically possible vertex that lies peripheral to the observed data (Gerber 2019). These ancestral nodes that lie outside the multidimensional convex hull defined by the tip data can inflate disparity relative to tip-only disparity (Brusatte et al. 2011). Similarly, authors have previously noted that inclusion of ancestral nodes can inflate disparity by (nearly) doubling the number of data points included in the trait space (Brusatte et al. 2011), raising questions about whether such increases reflect genuine biological signal. However, our findings demonstrate that the trait space expansion from the inclusion of ancestors, particularly when fossil sampling density is high, does converge on the true disparity of the clade under pre-ordination estimation. As Figure 4 shows, underestimation of disparity is ubiquitous to all discrete character estimation methods, which has previously been linked to character exhaustion (Erwin 2007). Ultimately, pre-ordination discrete estimations are less susceptible to extreme underestimation because ancestral nodes are not bound to a weighted average of the tip data in the way post-ordination estimation is.

### 4.2 Treatment of uncertainty

The choice of estimation method for continuous characters was largely driven by the treatment of uncertainty. At high fossil sampling densities, incorporating uncertainty through the use of distributions performed better than point estimates (Figure 2; Figure 3). This is due to the tendency for point estimates to smooth over evolutionary history (Boyko and O’Meara 2024; Boyko et al. 2026), underestimating the morphological variation of ancestral taxa. However, in contrast to discrete traits, continuous estimation distributions are unbounded and thus sensitive to data quality. Therefore, the variation introduced from sampling across a distribution was dependent on the density of fossils; at high fossil density, confidence intervals are constrained due to lower uncertainty in estimates. As fossil sampling decreases, these confidence intervals are widened and more noise is introduced as a result of using distribution (Schluter et al. 1997; Martins 1999), negating the advantage observed at higher fossil sampling.

For discrete characters, we found an inverse pattern where distribution methods were optimal at low fossil sampling densities and point estimates were optimal at high fossil sampling densities. This is likely due to the bounded nature of discrete traits; even with minimal information, the disparity error that can be returned is bounded by the number of states present at each node, which contrasts the effectively infinite error that can be returned by continuous characters. Regarding pre-ordination estimations, our results reveal two issues relating to uncertainty. At low fossil sampling or slow transition rates, the predominant issue facing the recovery of a true trait space is the lack of data. For fossil sampling, the lack of evidence in deeper time means our ancestral state estimates, and subsequent trait space pattern, are biased by the living tip data. In the case of slow transition rates, we face a paradoxical situation with slow rates where the root state can be easily estimated, yet the few transitions make the rate of change hard to estimate (Gascuel and Steel 2020). Using the hypercube analogy again, the state scaled likelihoods at each node under slow rates or low/absent fossils will be biased against transitions. Therefore, because point estimations essentially recover the most parsimonious evolutionary history, ancestral nodes will be constrained within a certain region of vertices in the trait space.

In contrast, pre-ordination distribution estimations allow the exploration of other vertices by sampling across alternative discrete state combinations. In the case of slow rates and low fossil sampling, this was found to improve disparity estimations (Figure 5). The flip side of this is the issue of saturated data, where the paradox is reversed and instead, we have an abundance of information, thus transition rates are easy to estimate, yet due to the rate of change most nodes are left with inconclusive scaled likelihoods. Under the fast rates simulated in this study, character transitions are saturated, so there is very little phylogenetic signal. The use of distributions amplifies the noise in this scenario, thus point estimates can be used to filter out the noise which is why they outperform distributions at medium/fast rates (Figure 5).

### 4.3 Model misspecification and fossil inclusion

As expected, we found that the differences in true and simulated disparity across models of evolution reflect model misspecification in ancestral state estimation. The inclusion of ancestral state estimations recovered a more accurate trait space than excluding ancestral states across all fossil sampling densities and disparity metrics when Brownian motion was the evolutionary model. Ancestral state estimation of continuous characters uses the BM model as an assumption for the evolution of traits, so this was expected. The assumptions of BM include a fixed mean and trait variance increasing linearly through time (Felsenstein 1985; O’Meara et al. 2006). We found that when these assumptions of BM were violated, such as under BM + trend and OU models, disparity accuracy returned by methods with ancestral state estimation decreased. For example, under low/absent fossil sampling, methods of ancestral state estimation performed worse than bypassing ancestral state estimation for BM + trend. In this case, disparity estimates tended to be underestimated, because state estimations will be intermediates of the observed tip data, absent to the directional trend that occurred and thus missing past diversity. Our results suggest a threshold between 5-15% fossil sampling where ancestral state estimation becomes optimal over its exclusion in disparity (Figure 3). This finding supports previous work that highlights the critical impact of fossil inclusion when directional trends are present in the trait data (Finarelli and Flynn 2006; Albert et al. 2009; Slater et al. 2012; Finarelli and Goswami 2013; Gearty et al. 2024).

Similarly, calculations of disparity derived from OU simulations were also underestimated when using ancestral state estimations. The OU process is characterised by a stationary variance, which contrasts the BM assumption of an increase in variance through time. Therefore, because BM estimates internal nodes as a weighted average of the observed tip data, the ancestral nodes are compressed towards the centroid of the trait space. This explains why, when model misspecification is fundamentally incorrect, bypassing ancestral state estimation outperforms the use of ancestral state estimation, even with 100% fossil sampling. We therefore encourage model-fitting prior to performing ancestral state estimation to avoid model misspecification; several R packages allow model-fitting and estimation of continuous traits under an OU model (Revell 2012; Pennell et al. 2014; Clavel et al. 2015). Note that empirical studies of disparity tend to focus on clades that are suspected to have undergone major evolutionary shifts, such as adaptive radiations (Ruta et al. 2013; Stroud and Losos 2016) or mass extinctions (Bapst et al. 2012; Halliday and Goswami 2016; Liu et al. 2024), which are often associated with shifts in evolutionary rates or models. The effect of such shifts or heterogeneity in the context of the aims of this paper remains to be tested; previous work has found that rate changes are hard to detect and can lead to erroneous inferences in single-trait ancestral state estimations (King and Lee 2015; Chira and Thomas 2016; Harrington and Reeder 2017; but see Reyes et al. 2018). Several recent methodological developments have aimed to accommodate for rate heterogeneity in both continuous (Elliot and Mooers 2014; Smaers et al. 2016; Castiglione et al. 2020) and discrete ancestral state estimation (Boyko and Beaulieu 2021; Revell and Harmon 2024) and so the testing of these methods in the context of disparity is a potential future avenue of research.

### 4.4 Disparity metrics and recovering a true trait space

Previously, the choice of disparity metric has been probed as a critical step in the disparity pipeline, specifically for examining different aspects of trait space (Guillerme et al. 2020b, 2025). Indeed, our results exhibit significant differences in disparity error among metrics. For example, recovering the sum of quantiles was ultimately harder than recovering the sum of variances or mean pairwise distance when using discrete traits. However, the hierarchy of methods was relatively consistent across disparity metrics, suggesting that the relative performance of ancestral state estimation methods is robust to different disparity metrics. Consistent performance across multiple disparity metrics can reassure us that multiple aspects of the true trait space are recoverable, such as size and density. Where there is inconsistency in methods across metrics, we can infer that the true disparity value has been attained as a methodological artefact, rather than the recovery of a true trait space pattern. This can be observed in the post-ordination distribution estimation of discrete traits, which performed well under the sum of variances and mean pairwise distance metrics, but noticeably worse under sum of quantiles. From this we can infer that while the internal density of taxa was correctly recovered, the geometric constraint of post-ordination estimation ultimately limited its ability to recover true trait space expansion.

### 4.5 Conclusions and recommendations

Our findings present a convincing case for the use of ancestral state estimation in disparity analyses, a method that has often been neglected in disparity workflows. Furthermore, our results issue a stark reminder of the essential need for fossils during macroevolutionary workflows. The estimated proportion of fossils sampled across empirical datasets is typically less than 5% (Raup 1976), which ultimately limits our inferences into macroevolutionary patterns. However, in the case of discrete traits, the incorporation of ancestral states is more optimal than their exclusion across all fossil sampling densities, and the same is true of continuous traits when the correct model is used for estimation. Based on these findings, we can recommend the following guidelines for the use of ancestral state estimation in disparity analyses: 1) for continuous traits, perform model testing prior to running ancestral state estimation to infer the likely evolutionary model parameters, and then estimate ancestral states under this model, using the distributions of estimates in disparity analyses; 2) for fast/medium rate discrete traits, pre-ordination point estimates should be used when fossil sampling is suspected to be less than 15% and post-ordination distribution when fossil sampling is lower than this. Pre-ordination distribution should be used when transition rates are slow across all fossil sampling densities.

## Acknowledgments

CNS would like to thank Robin Beck, Russell Garwood and Vera Weisbecker for thoughtful discussions about the study. TG acknowledges funding from NERC-UKRI Grant NE/X016781/1.

## Author Contributions

**Conceptualisation** CN Scutt (CNS), GH Thomas (GHT), N Cooper (NC), T Guillerme (TG); **Data curation** CNS; **Formal analysis** CNS; **Funding acquisition** TG; **Investigation** CNS, TG; **Methodology** CNS, TG; **Project administration** CNS, GHT, NC, TG; **Resources** CNS, TG; **Software** CNS, TG; **Supervision** GHT, NC, TG; **Validation** CNS; **Visualisaton** CNS; **Writing - original draft** CNS; **Writing - review & editing** CNS, GHT, NC, TG.

## Data Availability

The code to reproduce the analyses is available here https://github.com/calebowski/acedisparity (Scutt and Guillerme 2026). Data is available at https://doi.org/10.5061/dryad.280gb5n4p.

## Competing Interests

We declare no conflict of interest.

## Notes

### Competing Interest Statement

The authors have declared no competing interest.

