## Supplementary for "Disparity analyses are robust to ancestral state estimation uncertainty"

### S1 Supplementary Tables

Available on Dryad at `https://doi.org/10.5061/dryad.280gb5n4p`.

**Table S1** `/continuous/continuous_aggregated_method_multcomp.csv`

**Table S2** `/continuous/continuous_aggregated_method_model_fossil_multcomp.csv`

**Table S3** `/discrete/discrete_aggregated_method_multcomp.csv`

**Table S4** `/discrete/discrete_aggregated_method_model_fossil_multcomp.csv`

### S2 Supplementary Figures

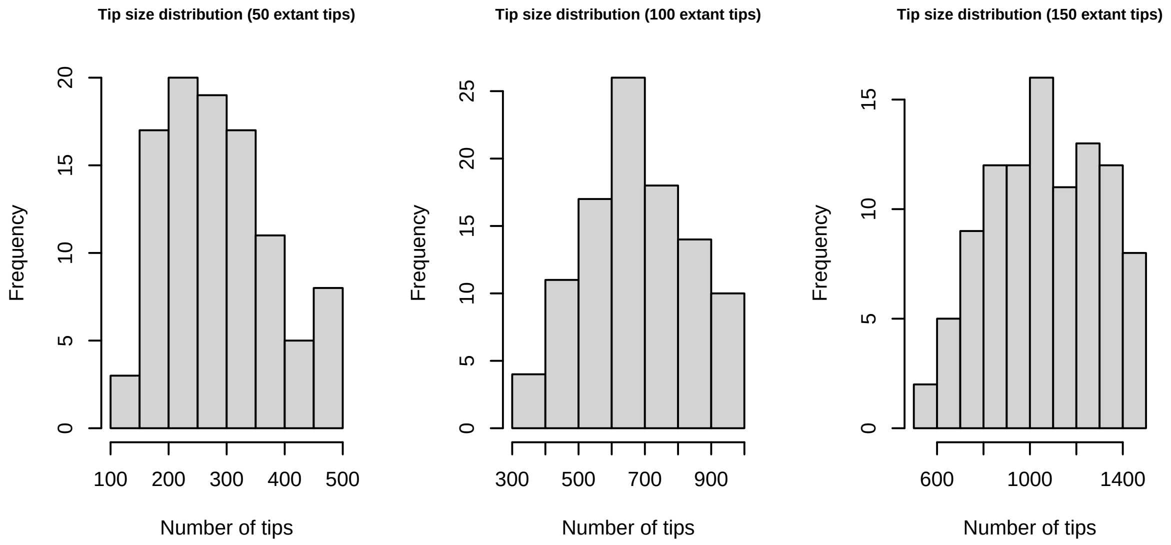


Figure S1: **Distribution of simulated tree sizes.** Histograms show the final number of extant tips across the three targeted size regimes (50, 100, and 150 extant taxa). Minor variations around the target sizes reflect the stochastic nature of the birth-death simulation process.

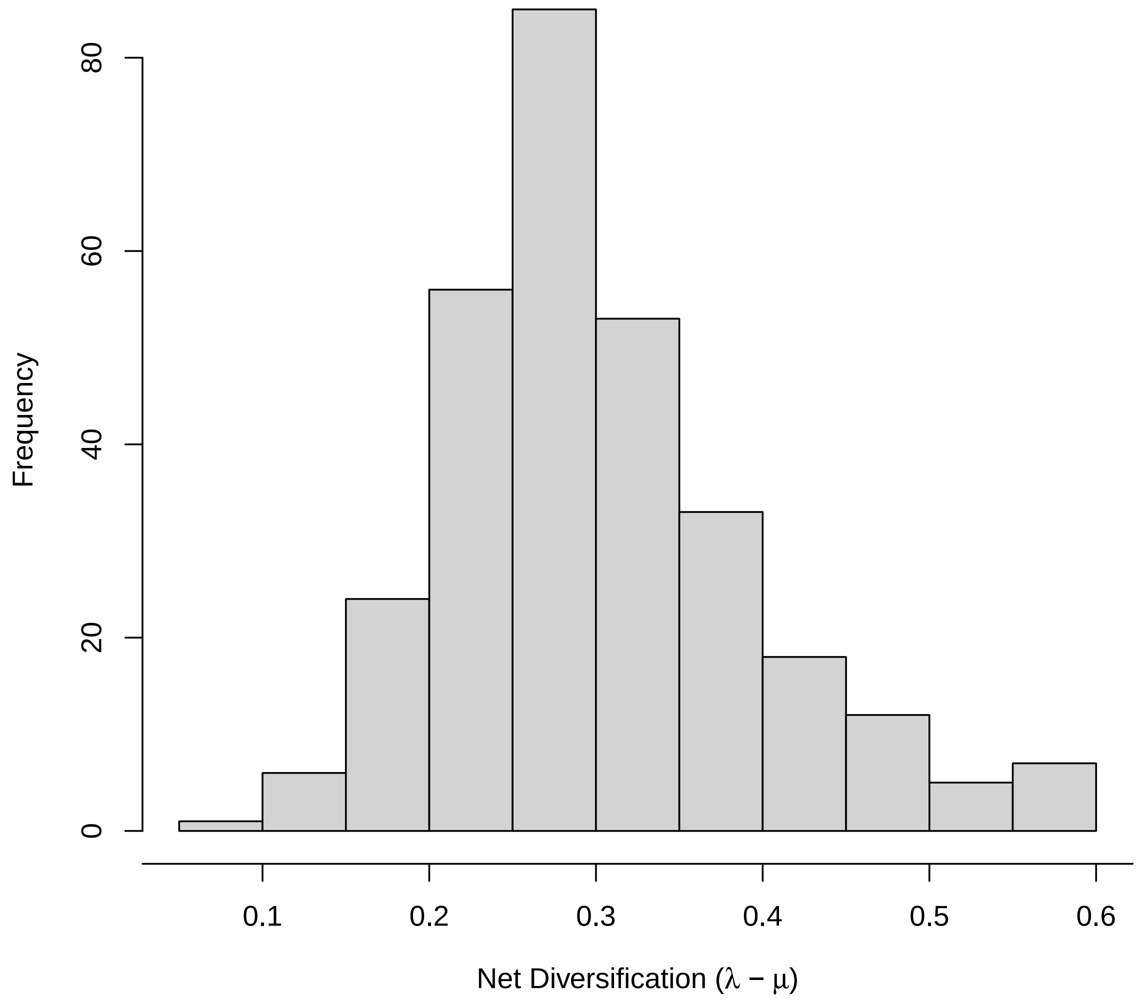


Figure S2: **Birth-death parameters for tree simulation.** The distribution of net diversification (speciation minus extinction) values across all simulated phylogenies.

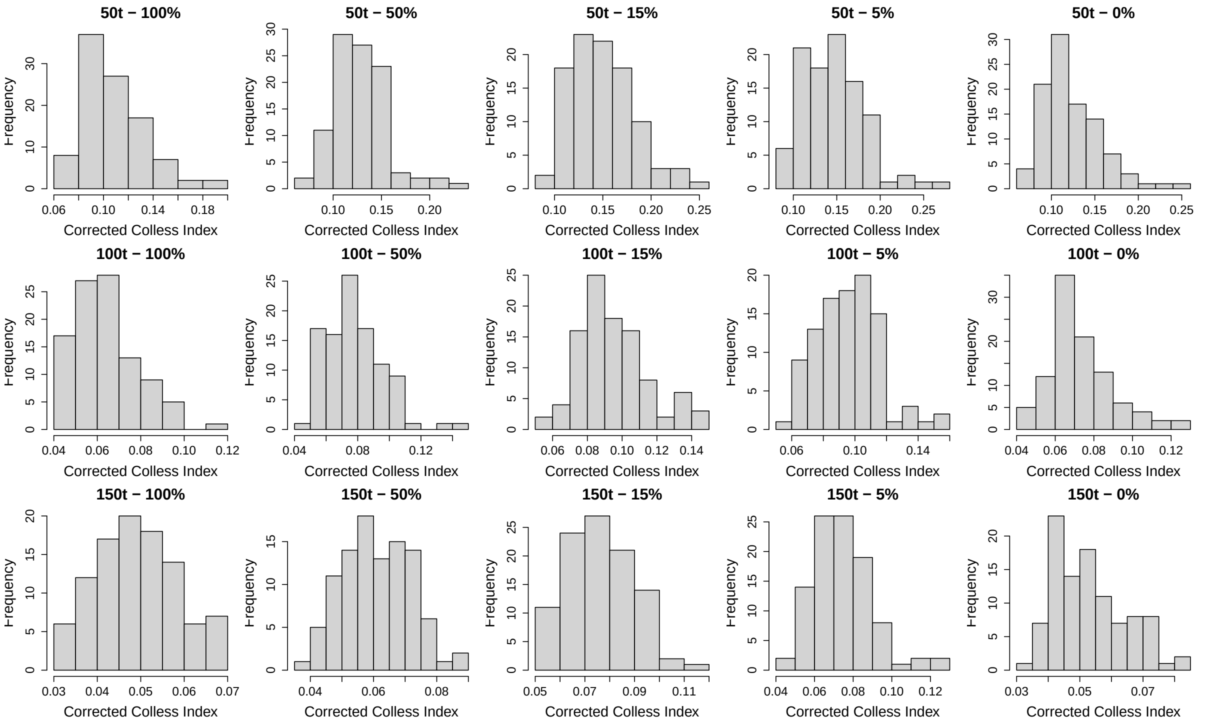


Figure S3: **Tree symmetry distributions.** Distribution of tree symmetry values measured using the corrected Colless index (Heard, 1992).

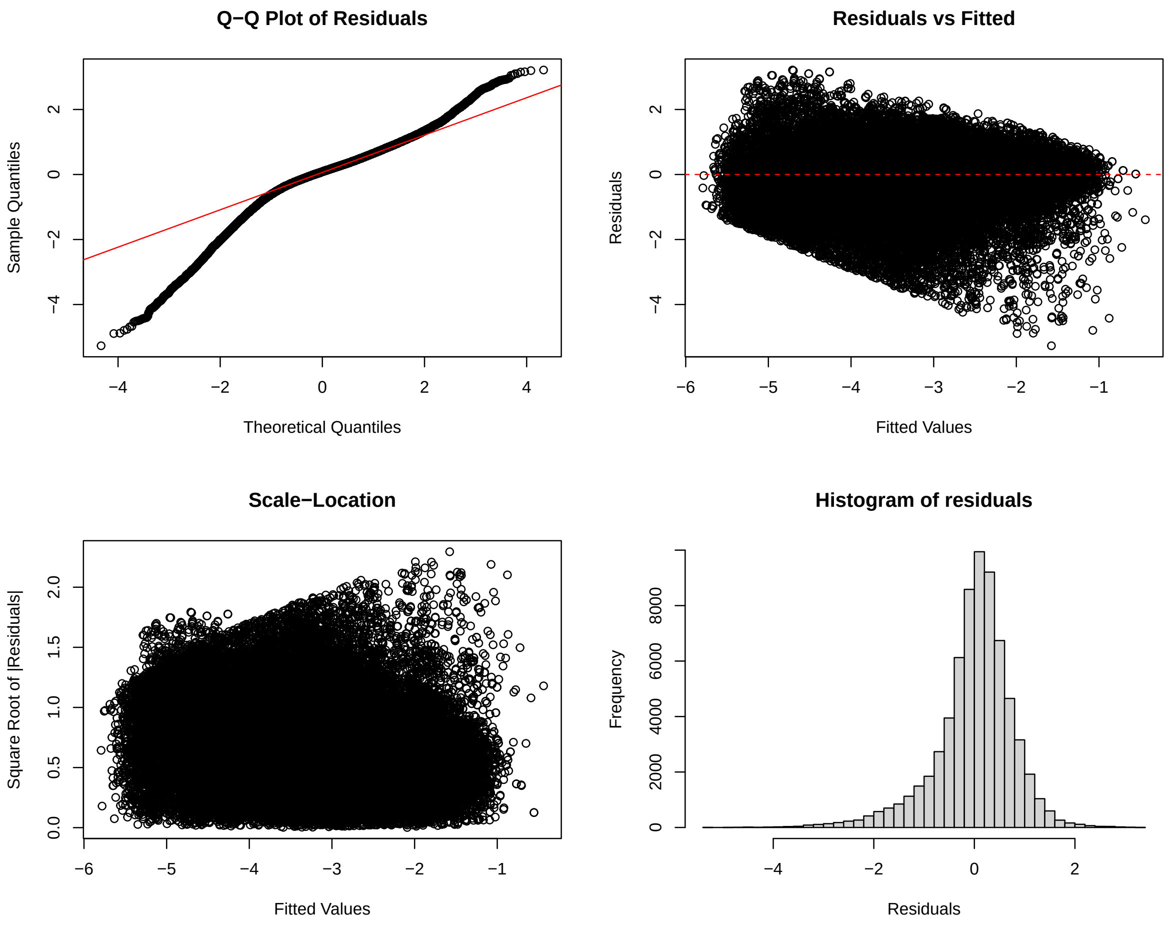


Figure S4: **Diagnostic plots for the continuous trait linear mixed-effects model (aggregated).**

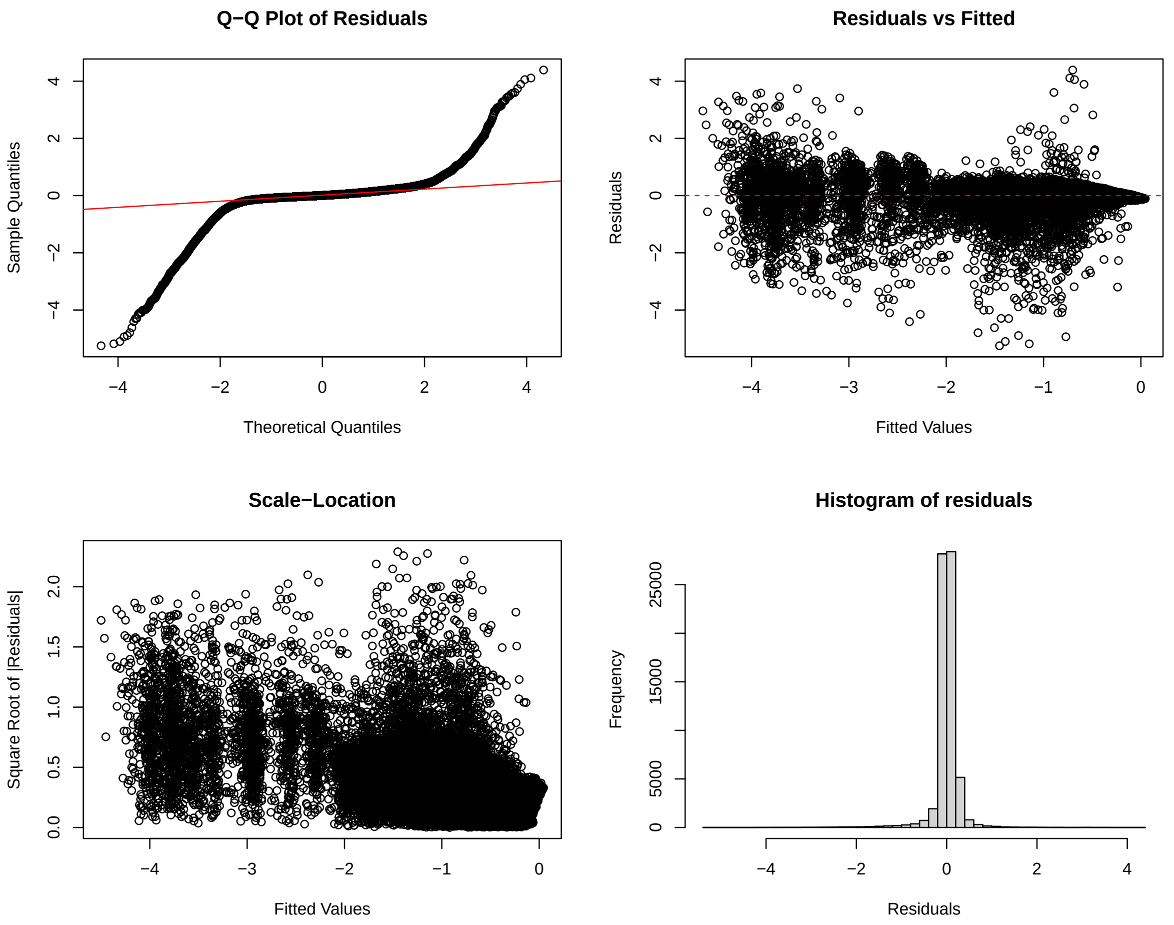


Figure S5: **Diagnostic plots for the discrete trait linear mixed-effects model (aggregated)**

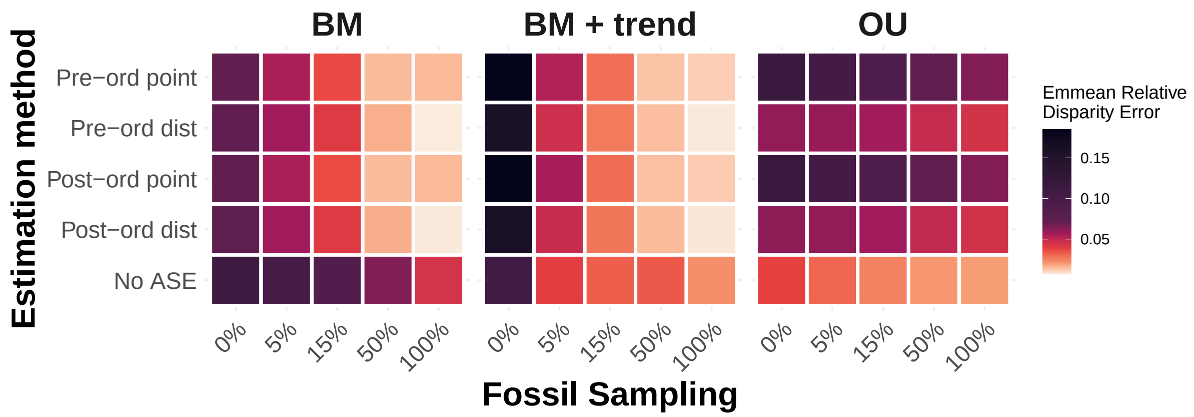


Figure S6: **Weighted LMM continuous heatmap**

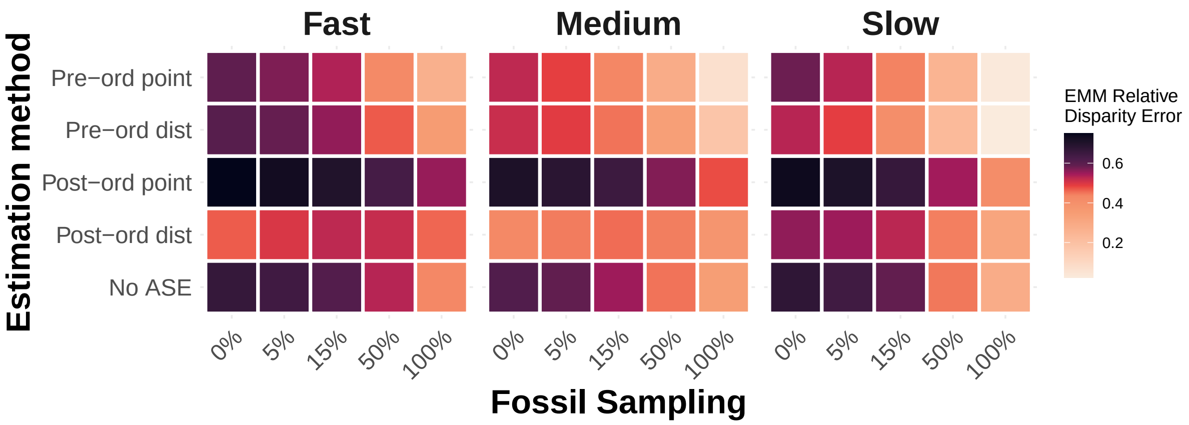


Figure S7: **Weighted LMM discrete heatmap**
